# The Serotonergic Neural Circuit Between The Dorsal Raphe Nucleus And The Basolateral Amygdala Is Implicated In Modulating The Arousal from Sevoflurane Anesthesia

**DOI:** 10.1101/2022.10.12.511924

**Authors:** Qian Yu, LeYuan Gu, XiTing Lian, YuLing Wang, Qing Xu, HaiXiang Ma, Lu Liu, WeiHui Shao, JiaXuan GU, Yue Shen, LiHai Chen, HongHai Zhang

## Abstract

Although some advancements concerning the arousal involved in mediating the delayed emergency from general anesthesia, which will lead to the serious complications, had been made, the role by arousal in modulating in delayed emergency still remains to be unclear. In our models, based on our previous working that activation of the 5-Hydroxytryptamine (5-HT) neurons in the dorsal raphe nucleus (DRN) by optogenetics can significantly reduce the emergency time by activating arousal pathway, we further test whether the serotonergic neural circuit between the DRN and the basolateral amygdala (BLA) is implicated in modulating the arousal from the sevoflurane anesthesia and the emergency time of sevoflurane anesthesia by the pharmacological, optogenetics and fiber photometry. Our findings showed that whether the serotonergic neural circuit between the DRN and the basolateral amygdala (BLA) plays a key role in modulating the arousal from the sevoflurane anesthesia and the emergency time of sevoflurane anesthesia. Based on the serotonergic neural circuit, the 5-HT 1 A receptor is of great significance to mediate the arousal and the emergency time of the sevoflurane anesthesia.

**Graphical Abstract:** Figure1

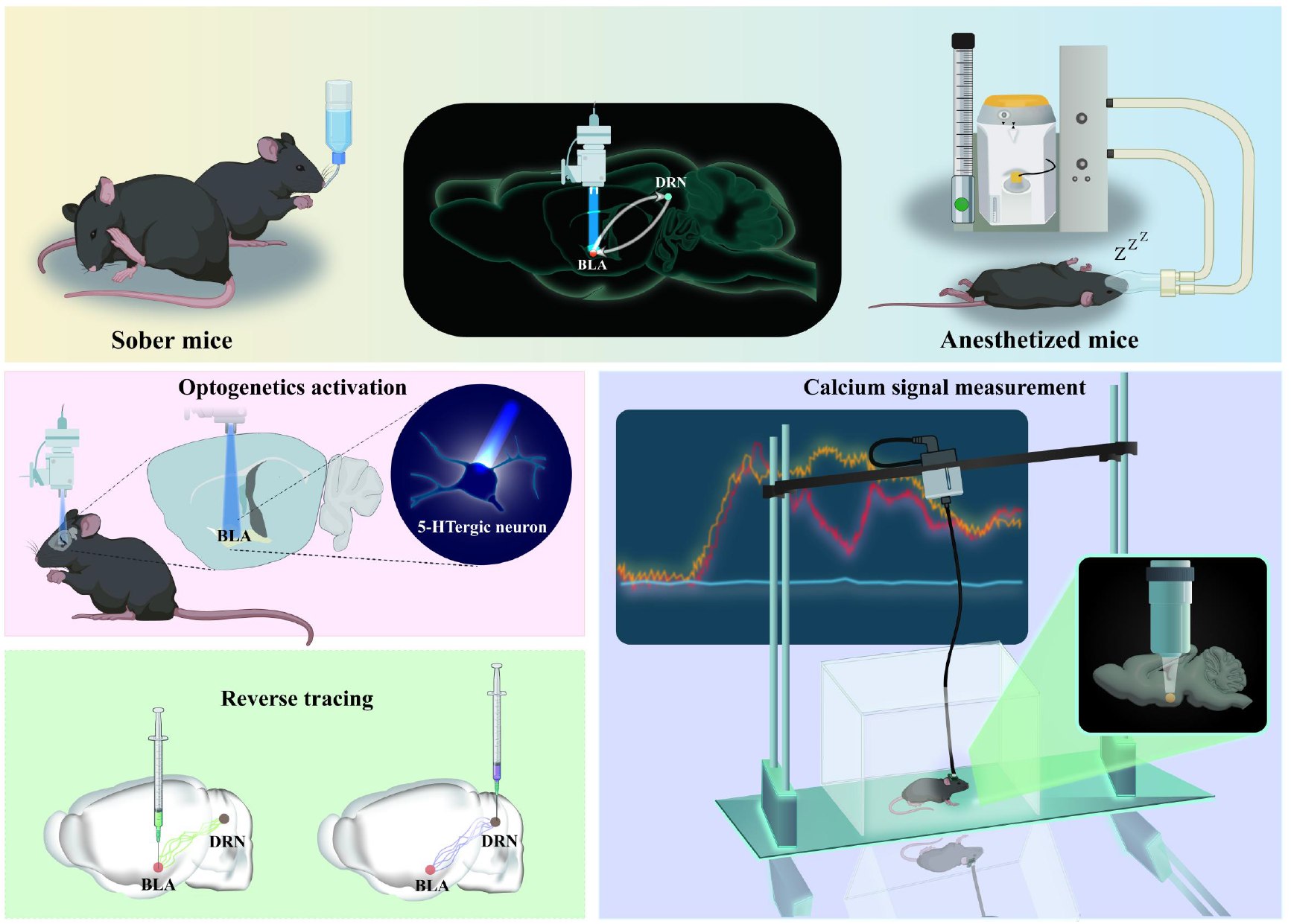

## Introduction

General anesthesia, which can be described as a pharmacologically induced state of amnesia, immobility, unconsciousness, and analgesia, has been widely used in surgery for nearly two centuries because of its ability to induce reversible loss of consciousness and maintain physiological stability^1^. General anesthesia is a reversible coma rather than analogous to any phase of sleep^2^. Previous studies have shown that under general anesthesia, the brain remains responsive to somatosensory, auditory, and visual stimuli, instead of silent, or unresponsive to external stimuli^3, 4, 5^. Most studies on the mechanism of arousal from general anesthesia in the past few years have focused on describing the action of anesthetic drugs on molecular targets^6^. Recently, researchers have begun to describe how anesthetics acted on specific neural circuits to cause the changes in arousal ^7, 8^, and shown that anesthesia-induced loss of consciousness occured through specific interactions with neural circuits in the central nervous system that regulated the endogenous sleep-wake system, such as the parbrachial nucleus, lateral hypothalamus, locus coeruleus, ventral tegmental area, ventrolateral preoptic nucleus^9, 10, 11, 12^, and optogenetic techniques were used in most of the experiments. However, whether other critical arousal promoting regions and their neural circuits are involved in the regulation of consciousness under general anesthesia is still unknown.

The amygdala plays an important role in initiating appropriate neurobehavioral responses to emotionally arousing events and activation of amygdala could promote REM and related activity. The amygdala is classified into three areas, as the basolateral, cortical, and central nuclei, and the functions are heterogeneous ^13^. Although many studies have not distinguished functional roles for nuclei of the amygdala, the available evidence suggests that the basolateral nucleus of the amygdala (BLA) is also involved in regulating REM and sends outputs to the central amygdala, sending neural information to the hypothalamus and the brainstem ^14, 15^. In behavioral control over breathing, the limbic cortex, especially the amygdala, is a major regulator of the emotion-associated breathing^16^.

It has been suggested that basal dopamine release from the BLA increases RR via local postsynaptic D2-like receptors, which might be related to the mechanisms of enhancement or inhibition of breathing^17^. It has been found that breathing and awakening are closely related. Just as sleep status can affect breathing control, changes in breathing can also lead to changes in wakefulness. Mice genetically deficient in serotonin neurons (Lmx1bf/f/p) in the central nervous system have been shown to have impaired arousal to CO_2_, but display normal arousal to hypoxia, sound, and mechanical stimuli, suggesting that serotonin neurons were involved in CO_2_ -induced sleep arousal^18^. 5-HT neurons, including midbrain neurons involved in sleep-wake regulation, respond to increased concentrations of inhaled CO_2_, resulting in increased firing rates, which may occur through a mechanism mediated by the 5-HT receptor^19, 20, 21, 22^. The dorsal raphe nucleus (DR), as the main source of 5-HT innervation in the forebrain, is involved in a variety of functions including anxiety, depression, and sleep-wake cycles, especially increased respiration and arousal induced by elevated levels of CO223. The electrical activity of its serotonergic neurons and the release of serotonin in the serotonergic projection area are increased during wakefulness and decreased during sleep. Furthermore, inactivation of the limbic structures that project to the hippocampus, such as pirifrom cortex, entorhinal cortex, or BLA, delay emergence from halothane and pentobarbital anesthesia and often reduced anesthetic-induced behavioral excitation^24, 25, 26^. Thus, we hypothesized a link between BLA and arousal from anesthesia and 5-HT in DR may also play a role in, but the precise neural mechanisms for this regulation, however, remain unclear.

## Methods

### 2.1 Animals

The experimental animals in this study are all wild type C57BL/6J mice, purchased from the Animal Experiment Center of Zhejiang University, and the experimental protocol was approved by the Experimental Animal Ethics Committee of Zhejiang University. All mice were housed in the barrier breeding room (SPF level), with appropriate temperature and humidity, clean living environment, and free access to food and water. For the experiments, eight-week-old mice weighing 20-25g were chosen. In order to avoid the interference of the estrous cycle of female mice, only male mice were used for behavioral experiments, which were completed from 9:00 to 15:00.

### 2.2 Sevoflurane anesthesia modeling and method of anesthesia induction and emergency time observation

The 8-week-old C57BL/6J mice were placed in a well-airtight inhalation anesthesia box (size, model size: length: 20cm, width: 10cm, height: 15cm; Reward) for 30 minutes prior to the experiment to adapt to the experimental environment. After 30 minutes of adaptation, the gas concentration of the sevoflurane volatilization tank was adjusted, and the anesthesia box was rotated 90 degrees every 15 s. When the mouse was in an abnormal position and could not turn itself prone onto all four limbs, the loss of righting reflex (LORR) was recorded as the Induction Time, and LORR was marked as the beginning of anesthesia maintenance phase. The corresponding experimental operation was carried out during maintenance phase in accordance with the experimental design. The sevoflurane volatilization tank was closed at the 30th minute of anesthesia to stop inhalation anesthesia, and the residual sevoflurane in the anesthesia box and pipeline was removed by rapid oxygenation, with the oxygen flow rate maintained at 2L/min. The anesthesia box was rotated 90 degrees every 15 s. The recovery of righting reflex (RORR) was considered if the mouse was in an abnormal position and could voluntarily place all four feet on the floor. The time between the end of anesthesia and RORR was recorded as the Emergency Time.

### 2.3 Stereotactic and virus microinjection surgery

8-week-old DBA/1 mice were anesthetized with 3.5% chloral hydrate and fixed in the head with a stereotactic apparatus (68018, RWD Life Sciences, Inc., Shenzhen, China). The body temperature of anesthetized mice were kept constant at 37°C during the procedure using a heating pad. For optogenetic viral delivery, pAAV-TPH2 PRO-ChETA-EYFP-WPRES-PAS (100nL, titer: 10^13^vg/ml, Shanghai Shengbo Biotechnology Company) or rAAV-TPH2-GCaMP6S-WPRE-hGH pA (100nL, titer: 10^12^vg/ml, Wuhan Zumi Brain Science and Technology Co., LTD.) was microinjected into unilateral or bilateral BLA (AP: -0.40mm; ML: + / - 2.95 mm; DV: -5.00 mm), based on mouse brain atlas (100nl, at a rate of 40 nl/min). Viruses were delivered via a gauge needle for the specification of 10ul controlled by an Ultra Micro Pump (160494 F10E, WPI). The syringe was not removed until 10 minutes after the end of the injection to allow the virus to spread. Then an optical fiber (FOC-W-1.25-200-0.37-3.0, Inper, Hangzhou, China) is implanted in the same area, and the optical fiber is located 0.05mm above the virus injection point (AP: -0.40mm; ML: + / - 2.95 mm; DV: 4.95 mm). Three weeks later, the next experiment was conducted. After the end of the experiment, the position of virus expression was confirmed, and the corresponding behavioral data was analyzed after observing whether the fluorescent expression of virus was in the target brain region.

Bilateral BLA cannula (O.D. 0.30 mm / 2.4 / M3.5 Arthur c., 62004, RWD life science co., LTD., shenzhen, China) was implanted the same as described above. The lateral ventricular cannula (O.D.0.41mm-27G/Pedestal 6mm/M3.5,62004, RWD Life Sciences Co., LTD., Shenzhen, China) was inserted as described previously with EEG electrodes embedded at the same time.

CTB-555 100nl (1μg/μl, BrainVTA Technology Co., LTD., Wuhan, China) was injected in DR (AP−4.55mm, ML−0.44mm, DV−2.80mm, 10° right) or left BLA(AP: -0.40mm; ML: + / - 2.95 mm; DV: -5.00mm) with a total content of 100nL for retrograde labeling of projection neurons, and then perfused after 1 week.

### 2.4 Pharmacological and optogenetics experiments

#### 2.4.1 Effect of intraperitoneal and BLA nuclear microinjection of 5-HTP on sevoflurane anesthesia induction time, emergency time and calcium signal of 5-HT neurons in BLA

5-HTP is the precursor of 5-hydroxytryptamine, which is soluble in normal saline. C57BL/6J mice were intraperitoneally injected with 5-HTP (50mg/kg, 100mg/kg), and injected again 24 hours later.One hour later, the mice were placed in the anesthesia box for induction and maintenance, and the induction time was recorded. Anesthesia was maintained for 30 minutes, and the emergency time was recorded then. The mouse in each group(vehicle, 50mg/kg, 100mg/kg) was subjected to three experiments, with complete metabolism of sevoflurane at an interval of one week.

One week after the implantation of bilateral cannulas in 8-week-old C57BL/6J mice, the BLA nuclear microinjection experiment was conducted. The mice were placed in the anesthesia box for anesthesia induction and the induction time was recorded. After 15 minutes of anesthesia, 200nl of 5-HTP (5mg/kg, 10mg/kg) was injected into the bilateral cannulas of BLA. Anesthesia was stopped at 30 minutes, and the emergency time was recorded. Each mouse in different group (vehicle, 5mg/kg, 10mg/kg) was subjected to three experiments, with complete metabolism of sevoflurane at an interval of one week.

#### 2.4.2 Effect of photostimulation of 5-HT neurons in BLA on induction time and emergency time of sevoflurane anesthesia

The experiment was performed in 8-week-old C57BL/6J mice 3 weeks after virus injection and 1 week after fiber implantation. The mice were placed in the anesthesia box for 30min before anesthesia induction and the induction time of anesthesia was recorded. At the 15th or 20th minute of anesthesia maintenance, the mice were exposed to blue light photostimulation (465 nm, 20 Hz, 20ms pulse, 10mW or 15mW) for 10 min or 15 min. After anesthesia maintenance for 30min, the anesthesia was stopped and the emergency time was recorded. The mouse in each group was subjected to three experiments, with complete metabolism of sevoflurane at an interval of one week.

#### 2.4.3 Effects of lateral ventricle injection 5-HT1A receptor antagonist and photostimulation of BLA 5-HT neurons on the sevoflurane anesthesia induction time, emergency time and electroencephalogram (EEG)

The experiment was performed in 8-week-old C57BL/6J mice 3 weeks after virus injection and 1 week after lateral ventricle cannula implantation. EEG recording was started 5 minutes prior to induction. After injecting 5-HT1A receptor antagonist WAY 100635 (10mg/ml) through the lateral ventricle cannula, the mice were put into the anesthesia box and the induction time was recorded. At the 15th minute of anesthesia maintenance, the mice were exposed to blue light photostimulation (465nm, 15mW, 20Hz, 20ms pulse) for 15min. After 30 minutes of anesthesia, anesthesia was stopped and the emergency time was recorded. After the recovery of righting reflex, EEG was recorded for another 10 minutes. The mouse in each group (vehicle+no light group, 1A (-)+no light group, vehicle+light group, 1A (-)+light group) was subjected to four experiments, with complete metabolism of sevoflurane at an interval of one week.1A (-)= WAY 100635

### 2.5 Fiber photometry

#### 2.5.1 Calcium signal changes of 5-HT neurons in BLA at different stages under sevoflurane anesthesia

The experiment was performed in 8-week-old C57BL/6J mice 3 weeks after virus injection and 1 week after optical fiber implantation. The mice were placed in the anesthesia box and recorded calcium signal 5min prior to induction when they were awake. Then the anesthesia induction was conducted and maintenance lasted for 30min. The induction time and emergency time was recorded at the same time. After the recovery of righting reflex, the experiment was stopped after 10min of optical fiber recording.

#### 2.5.2 Calcium signal changes of 5-HT neurons in BLA under sevoflurane anesthesia after intraperitoneal injection of 5-HTP

The experiment was performed in 8-week-old C57BL/6J mice 3 weeks after virus injection and 1 week after optical fiber implantation. C57BL/6J mice were intraperitoneally injected with 5-HTP (50mg/kg, 100mg/kg), and injected again 24 hours later. One hour later, the mice were placed in the anesthesia box for induction and maintenance, and the induction time was recorded. The recording of calcium signal was started 5min prior to induction when they were awake. After 30 minutes of anesthesia, the anesthesia was stopped and the emergency time was recorded. After the recovery of righting reflex, the experiment was stopped after 10min of optical fiber recording. The mouse in each group was subjected to experiments twice (vehicle group, 10mg/kg group) with one week interval between each experiment to completely metabolize sevoflurane.

#### 2.5.3 Changes of calcium signal and c-fos activation of 5-HT neurons in BLA under sevoflurane anesthesia after blue light activation of DRN

The experiment was performed in 8-week-old C57BL/6J mice 3 weeks after virus injection and 1 week after optical fiber implantation. The mice were placed in the anesthesia box and recorded for 5min prior to induction when they were awake. Then the anesthesia induction was conducted and the induction time was recorded at the same time. At 15min of maintenance, the mice received blue light photostimulation (465nm, 15mW, 20Hz, 20ms pulse) in DRN for 15 min. After 30min maintenance, the anesthesia was stopped and the emergency time was recorded. After the recovery of righting reflex, the optical fiber recording was continued for 10min.

### 2.6 Immunohistochemistry

The placement of the optical fiber cannula tip including intracerebroventricular (ICV) and microinjection of 5-HTP with bilateral BLA in each implanted mouse was verified by histology. After the experiment, C57BL/6J mice were sacrificed and perfused with PBS and 4% PFA. After saturated in 30% sucrose (24h), each brain was sectioned into 35μm thickness of coronal slices with a freezing microtome (CM30503, Leica Biosystems, BuffaloGrove, IL, USA), the sections were first washed in PBS 5mins for 3 times and then incubated in blocking solution containing 10% normal donkey serum (017-000-121, Jackson ImmunoResearch, West Grove, PA), 1% bovine serum albumen (A2153, Sigma-Aldrich, St. Louis, MO), 0.3% Triton X-100 in PBS for 2 h at room temperature. Then, for c-fos or TPH2 staining in DR or BLA, sections were incubated at 4°C overnight in rabbit anti-c-fos(1:1000 dilution, 2250T Rabbit mAb, Cell Signaling Technolog, Danvers, Massachusetts, USA) primary antibody or mouse anti-TPH2 (1:100 dilution, T0678, Sigma-Aldrich, St. Louis, MO) primary antibody and donkey anti-mouse Alexa 546 (1:1000; A10036, Thermo Fisher Scientific, Waltham, MA, USA) secondary antibody or goat anti-rabbit cy5 (1:1000; A10523, Thermo Fisher Scientific, Waltham, MA, USA) secondary antibody for 2h at room temperature. After washing with PBS 15mins for 3 times, the sections were mounted onto glass slides and incubated in DAPI (1:4000;Cat# C1002; Beyotime Biotechnology; Shanghai, China) 7mins at room temperature.Finally, the glass slides were sealed sheet by Anti-fluorescence attenuating tablet.All images were taken with Nikon A1 laser-scanning confocal microscope (Nikon, Tokyo, Japan).The numbers of immunopositive cells were counted and analyzed using ImageJ (NIH, Baltimore, MD). It is worth noting that the mouse of the implantation placement out of the target of brain structure will be abandoned in our experiments. The positive cells co-expression c-fos, and TPH2 were counted as previously described.

### 2.7 Data statistics and analysis

The experimental data were reported as mean ± SD. Shapiro-wilk normality test was performed for all experimental data before data analysis. Comparison between the two groups: If the data conformed to the normal distribution, Student t-test was used, including independent sample T-test and paired sample T-test. If the data did not meet the normal distribution, Mann-Whitney U or Wilcoxon signed-rank test was used. Levene test was used to detect the homogeneity of square difference. After the data met the normal distribution and homogeneity of variance, one-way ANOVA analysis was used for the comparison of three or more groups, and two-way ANOVA analysis and Bonferroni’s test were used for the comparison between two factors and different groups. Statistical significance was inferred if p < 0.05. GraphPad Prism TM 8.0 and SPSS version 26.0 were used for statistical analysis of data in this study.

## Results

### 3.1 Sevoflurane anesthesia modeling of C57BL/6J mice

The time to LORR of mice exposed to different sevoflurane concentrations was different (Fig 1B). In order to determine the appropriate concentration of sevoflurane, we normalized the Induce time of mice in each group, and took the drug concentration corresponding to the median Induce time of all mice, i.e. 2.55% sevoflurane, as the induction and maintenance dosage for subsequent mice (Fig 1).

**Figure1.**
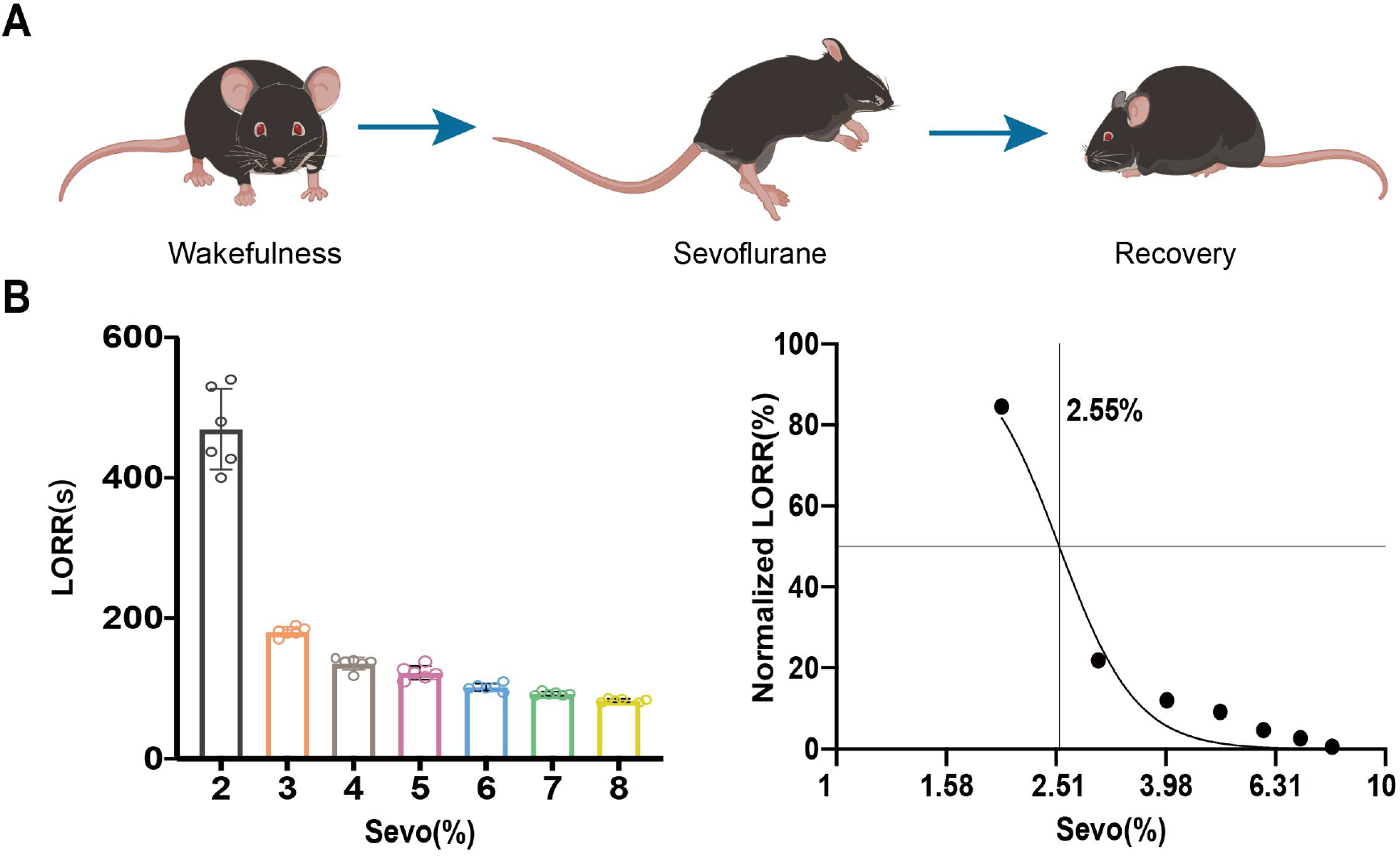
Sevoflurane anesthesia modeling of C57BL/6J mice. **A**. The anesthesia process of C57BL/6J mice **B**. The time to LORR under different inhalation concentrations of sevoflurane (n=6 for each group). **C**. Normalization the Induce time of mice in each group. **LORR, loss of righting reflex; Sevo, sevoflurane**

### 3.2 5-HT neuronal activity in BLA during maintenance period was significantly reduced based on photometry recordings

During sevoflurane anesthesia, calcium signals of 5-HT neurons in bilateral BLA infected with GCaMP6s were recorded by photometry (Fig 2). There was significant difference in peak ΔF/F of 5-HT neurons in BLA between the maintenance period and other stages (p<0.05, Fig2). These data indicates that the activity of 5-HT neuron in BLA may involve in the process of sevoflurane anesthesia.

**Figure2.**
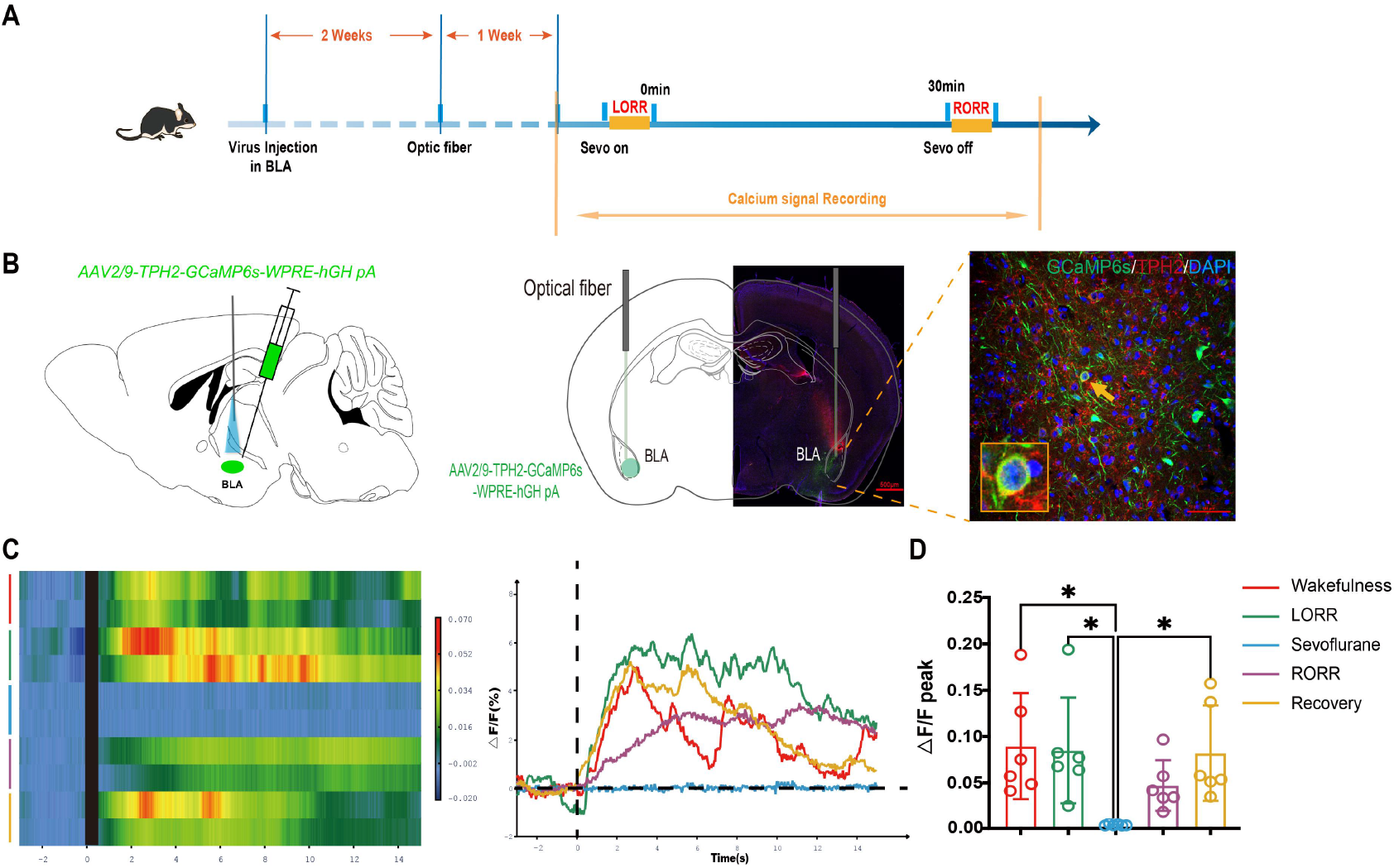
Changes of calcium signaling in BLA 5-HT neurons during anesthesia. **A**. Schematic illustration of Photometric recordings **B**. A representative photomicrograph shows microinjection and optical fiber locations and the co-expression GCaMP6s and TPH2. **C**. The heatmap and statistical diagram of calcium signaling changes in bilateral BLA 5-HT neurons during sevoflurane anesthesia. **D**. Peak ΔF/F in BLA during the maintenance of anesthesia was significantly lower than that during the waking, induction and recovery periods. **\*p<0.05**

### 3.3 5-HT neuronal activity in BLA during anesthesia was significantly increased by elevating the level of 5-HT

Compared with the vehicle group, intraperitoneal injection of 5-HTP with different concentrations had no significant difference in Induction time (p>0.05, Fig 3). The group treated with 5-HTP 100mg/kg had longer Induce time than 50mg/kg (p<0.05, Fig 3). Compared with the vehicle group, the emergency time was significantly reduced after IP 100mg/kg 5-HTP (p<0.001). And compared with the group IP 100mg/kg 5-HTP, the emergency time was significantly increased after IP delivery of 5-HTP at a dosage of 50mg/kg (p<0.01, Fig 3).

**Figure 3.**
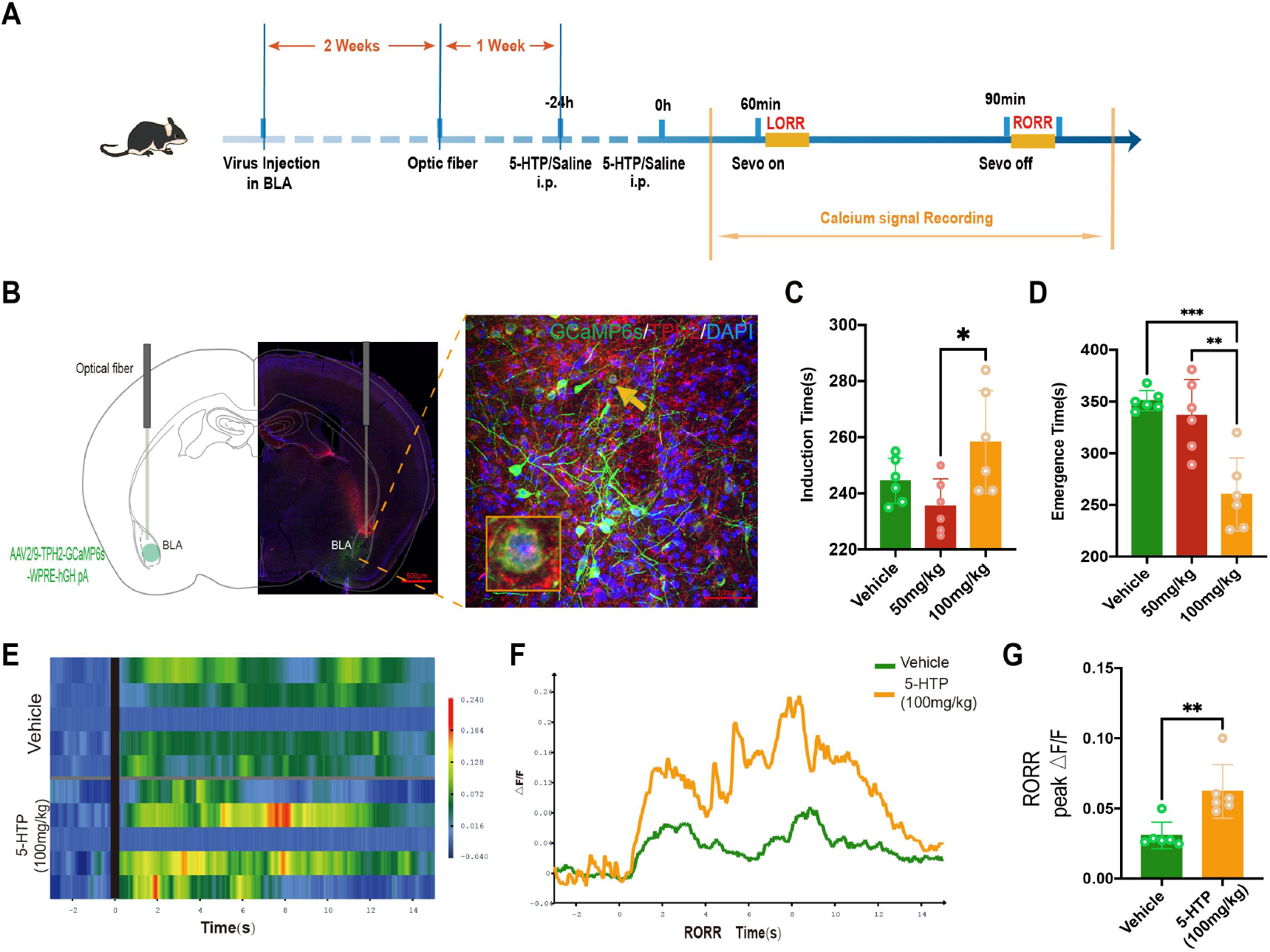
Effects of administration 5-HTP to elevate the level of 5-HT on the Changes of calcium signaling in BLA 5-HT neurons during anesthesia. A. Schematic illustration of Photometric recordings **B**. A representative photomicrograph shows microinjection and optical fiber locations and the co-expression GCaMP6s and TPH2. **C-D**. Compared with the vehicle group, intraperitoneal injection of 5-HTP with different concentrations had no significant difference in Induction time (p>0.05). The group treated with 5-HTP 100mg/kg had the longer Induce time than 50mg/kg (p<0.05). **E-F**. The heatmap and statistical diagram of calcium signaling changes in bilateral BLA 5-HT neurons during sevoflurane anesthesia with the treatment of 5-HTP and without the treatment of 5-HTP. **D**. Peak ΔF/F in BLA during the maintenance of anesthesia without the treatment of 5-HTP was significantly lower than that during the waking in the group with 5-HTP, induction and recovery periods. **\*p<0.05**

To further explore the role of 5-HT neurons in BLA, we microinjected 400nL vehicle and 5-HTP in BLA. Compared with the vehicle group, microinjection 5-HTP in BLA had no significant difference in Induction time (p>0.05, Fig 4 E). However, compared with the vehicle group, the emergency time was significantly reduced after microinjection 10mg/ml 5-HTP (p<0.001). And compared with the group microinjection 10mg/ml 5-HTP, the emergency time was significantly increased after microinjection 5-HTP at a dosage of 5mg/ml (p<0.01, Fig 4).

**Figure 4.**
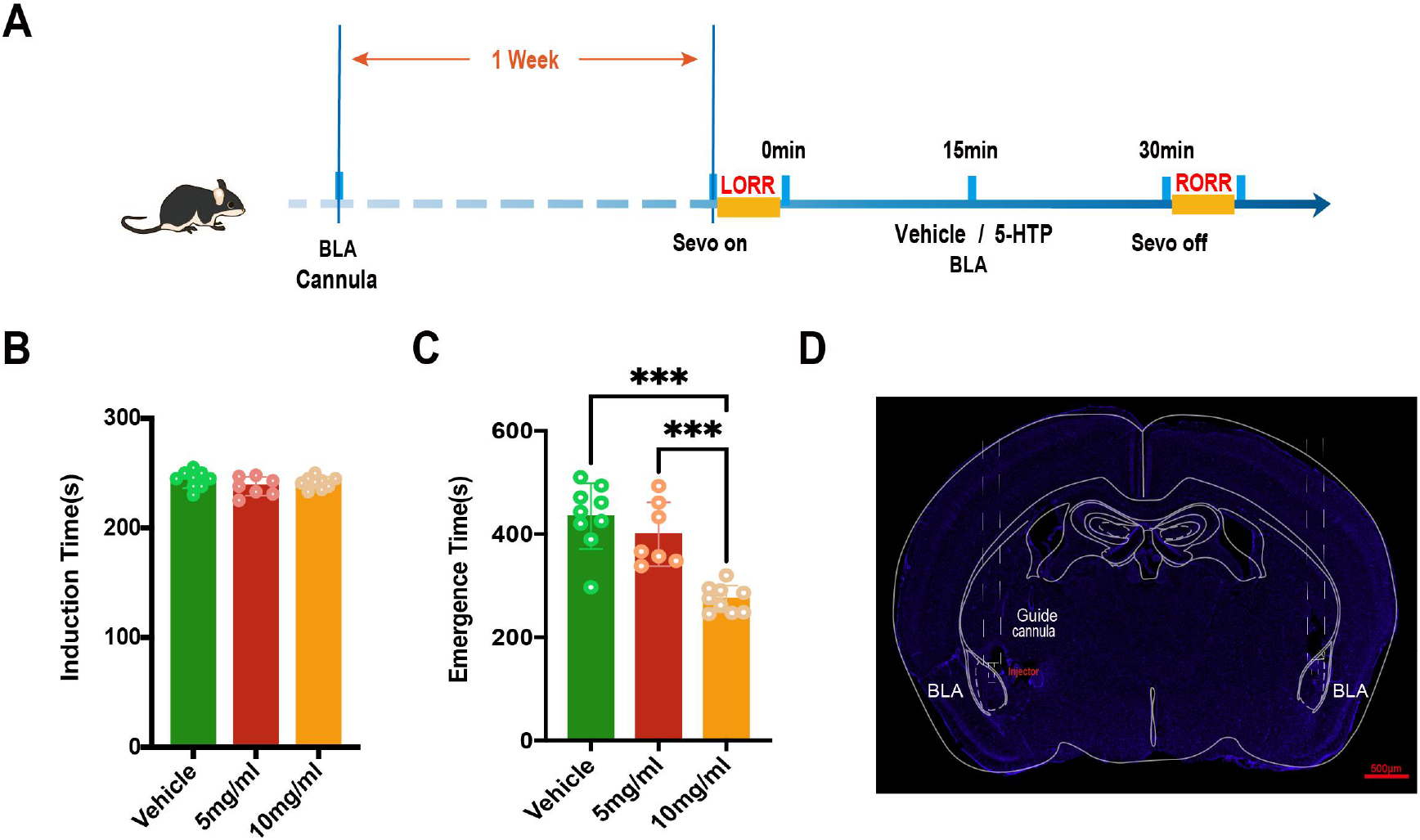
The difference of Induce time and Emergency time when elevating the level of 5-HT via IP and BLA microinjection 5-HTP. **A**. Schematic illustration of microinjection 5-HTP in BLA **B-C**. Effect of IP different concentrations of 5-HTP on Induction time and Emergency time **D**. A representative photomicrograph shows microinjection locations. **E-F**. Effect of microinjection different concentrations of 5-HTP in BLA on Induction time and Emergency time **G**. Peak ΔF/F of 5-HT neurons in BLA during RORR was significantly increased in the group treated with IP 100mg/kg 5-HTP. **\*\*\*p<0.001, **p<0.01, *p<0.05, IP intraperitoneal injection, RORR recovery of righting reflex**

There was significant difference in peak ΔF/F of 5-HT neurons in BLA between the vehicle group and the group IP 100mg/kg 5-HTP during RORR (p<0.01, Fig 3G-H). These data indicates that 5-HT neuron in BLA may have an important impact on the process of sevoflurane anesthesia, especially the recovery stage.

### 3.4 Effect of activation 5-HT neurons in BLA on Induction time and Emergency time of sevoflurane anesthesia

We examined the effect of selective enhancement of 5-HT neurotransmission on Induce time and Emergency time by applying photostimulation (blue light: 10min, 15mW; 15min, 15mW; 15min, 10mW) to 5-HT neurons in BLA. Compared with the vehicle group, photostimulation of the BLA (10min, 15mW; 15min, 15mW; 15min, 10mW) had no significant difference in Induction time (p>0.05, Fig 5 B). The group treated with 5-HTP 100mg/kg had the longer Induce time than 50mg/kg (p>0.05, Fig 5 C and 5 F). Compared with the vehicle group, the Emergency time was significantly reduced after photostimulation of the BLA (blue light, 15min, 15mW) (p<0.001). And compared with the group photostimulation of the BLA (blue light, 15min, 15mW), the Emergency time was significantly increased after photostimulation of the BLA (blue light: 10min, 15mW; 15min, 10mW) (p<0.05, Fig 4D; p<0.01, Fig 5 G). There was no significance in Induce time in the following group respectively: no PS, left PS, right PS and bilateral PS (p>0.05, Fig 5 E). And compared with the no PS group, the Emergency time was significantly decreased in the group with left PS, right PS or bilateral PS (p<0.001, Fig 5 H), indicating activation of 5-HT neuron located in BLA could promote the recovery of anesthesia in C57BL/6J mice.

**Figure 5.**
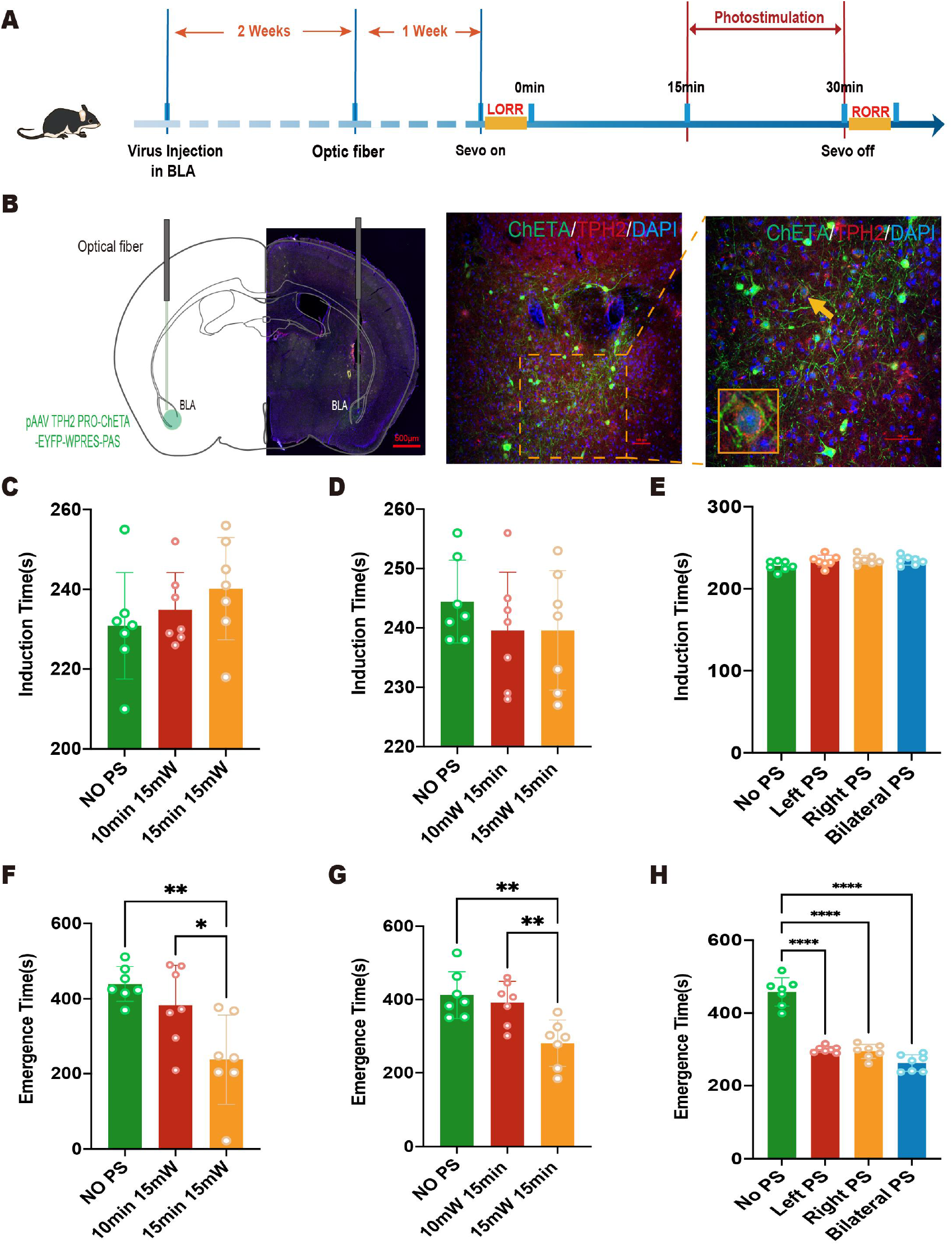
The difference of Induce time and Emergency time when photostimulating 5-HT neuron in BLA with different light parameters. **A**. Schematic illustration of photostimulation 5-HT neuron with different light parameters in BLA **B**. A representative photomicrograph shows the locations of optical fiber and virus expression. **C-D, F-G**. Effect of photostimulating 5-HT neuron in BLA (10min, 15mW; 15min, 15mW; 15min, 10mW) on Induction time and Emergency time **E, H**. Effect of photostimulating 5-HT neuron in BLA (no PS, left PS, right PS, bilateral PS) on Induction time and Emergency time. **\*\*\*p<0.001, **p<0.01, *p<0.05, PS photostimulation**

### 3.5 EEG activity upon optogenetic activation of 5-HT neurons in BLA-mediated reduction of the Emergency time during sevoflurane anesthesia

pAAV-TPH2 PRO-ChETA-EYFP-WPRES-PAS was delivered into BLA 3 weeks before anesthesia, and an optical fiber with a headstage for EEG was implanted 1 week before experiments. The mice were divided into three groups: no PS, PS (10mW, 15min), PS (15mW, 15min). The activity of cortical EEG of three groups decreased significantly after LORR, and kept at a low level during the maintenance of anesthesia. During the recovery stage, the cortical EEG recovered, but was still lower than that before anesthesia (Fig 6 A).

**Figure 6.**
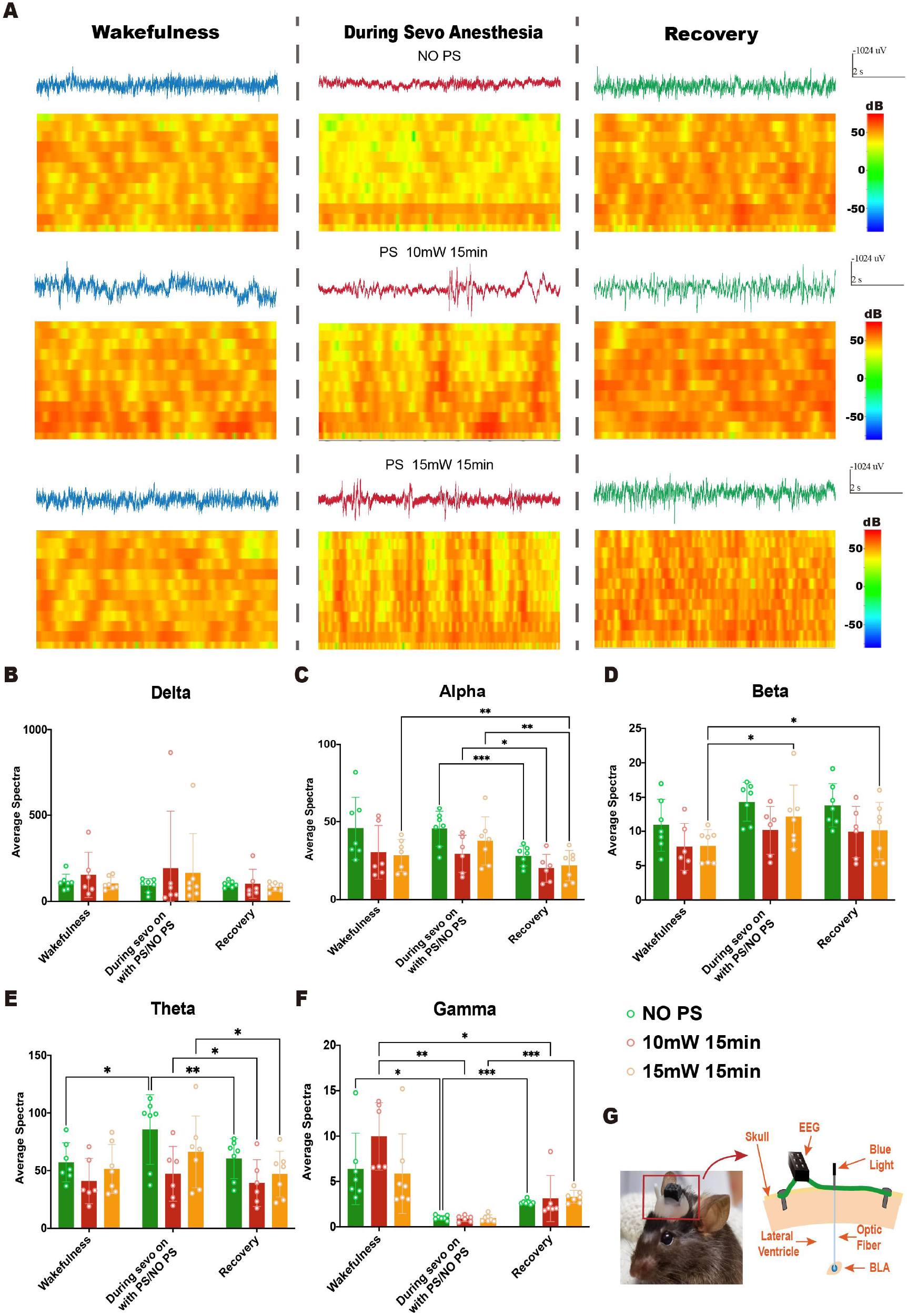
Effect of photogenetic activation of 5-HT neurons in BLA on EEG activity. **A**. EEG and spectrum of mice in vehicle group, PS (10mW, 15min) and PS (15mW, 15min) under awake, anesthetized and recovery states. **B-F**. Delta, Alpha, Beta, Theta and Gamma wave proportion of EEG in three groups of mice in different states **G**. Schematic Diagram of Mouse Cortical EEG Recording and optical fiber. **\*\*\* p<0.001, ** p<0.01, * p<0.05, PS photostimulation**

For Delta wave, the average proportion in the three groups during anesthesia had no significant difference (p>0.05, Fig 6 B). As for Alpha wave, the average proportion of Alpha wave in maintenance phase was significantly higher than that in the recovery phase in the vehicle group (P<0.001), PS (10mW, 15min) (P<0.05) and PS (15mW, 15min) (P<0.01). And the average proportion of Alpha wave in PS (15mW, 15min) during wakening phase was significantly higher than that in the recovery phase (P<0.01, Fig 6 C). For Beta wave, the group treated with PS (15mW, 15min) in the awake state was significantly lower than that in the anesthesia or recovery state (P<0.05, P<0.05, Figure 5 D). For Theta wave, the percentage of Theta wave in vehicle group during maintenance phase was significantly higher than wakefulness phase (P<0.05) and recovery phase (P<0.05). And the percentage of Theta wave in PS (10mW, 15min) and PS (15mW, 15min) during maintenance phase was significantly higher than recovery phase (P<0.05, P<0.05, Fig 6 E).

As for Gamma wave, there was no statistical difference in PS (10mW, 15min) during anesthesia. In the vehicle group, the Gamma wave proportion was significantly lower in the maintenance stage than that in wakefulness (P<0.05) and recovery (P<0.001). In the group with PS (10mW, 15min), the Gamma wave proportion was significantly higher in the wakefulness than that in maintenance (P<0.01) and recovery (P<0.05). In the group with PS (15mW, 15min), the Gamma wave proportion was significantly higher in the recovery than that in maintenance (P<0.001, Fig 6 F). These indicates EEG activity alters in different stages and different wave change in different ways and degrees.

### 3.6 EEG activity upon optogenetic activation of 5-HT neurons in BLA-mediated reduction of the Emergency time during sevoflurane anesthesia

pAAV-TPH2 PRO-ChETA-EYFP-WPRES-PAS was delivered into BLA 3 weeks before anesthesia, and an optical fiber with a headstage for EEG was implanted 1 week before experiments. The mice were divided into three groups: no PS, PS (15mW, 10min), PS (15mW, 15min). The activity of cortical EEG of three groups decreased significantly after LORR, and kept at a low level during the maintenance of anesthesia (Fig 7 A).

**Figure 7.**
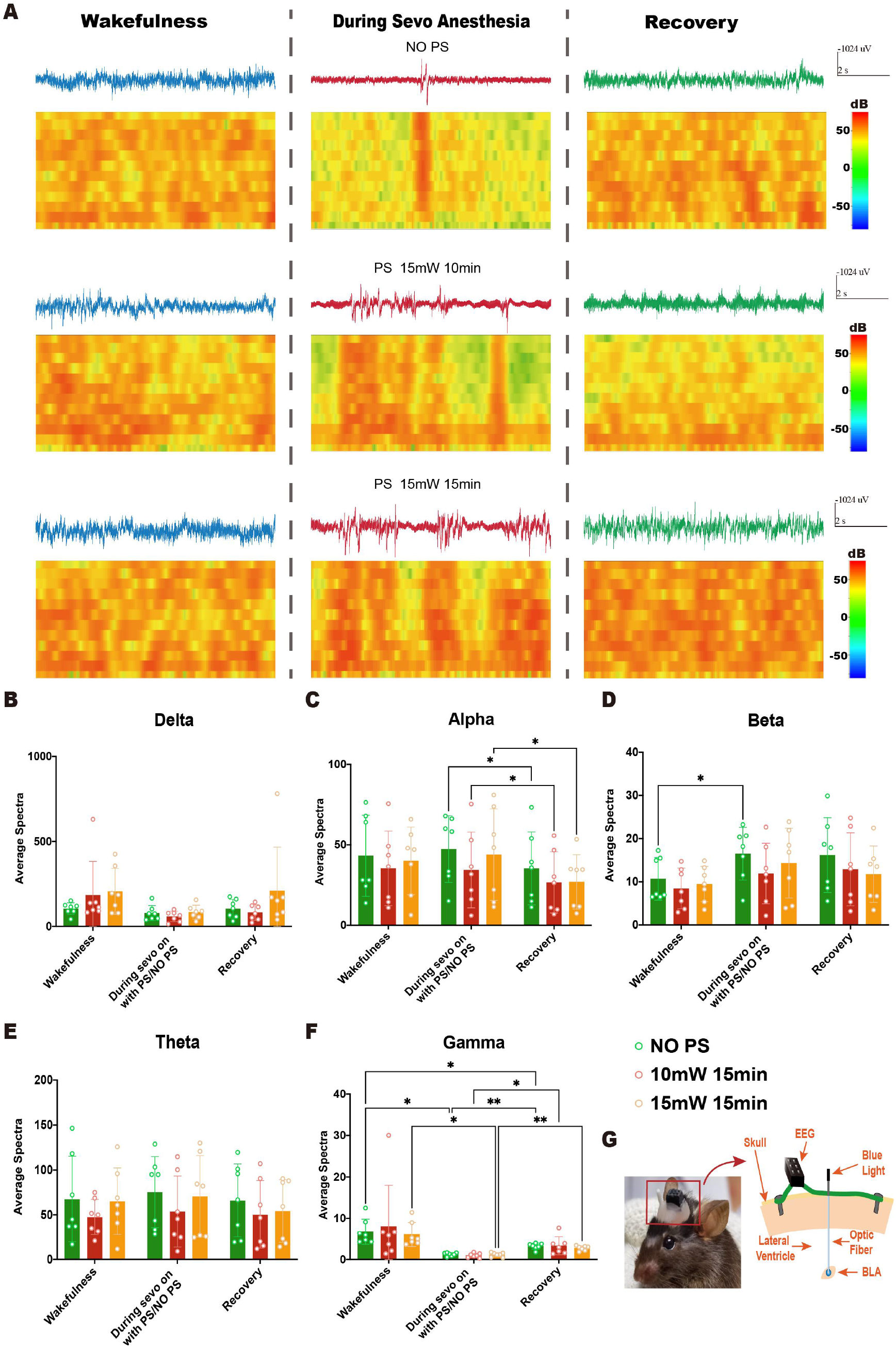
Effect of photogenetic activation of 5-HT neurons in BLA on EEG activity. **A**. EEG and spectrum of mice in vehicle group, PS (15mW, 10min) and PS (15mW, 15min) under awake, anesthetized and recovery states. **B-F**. Delta, Alpha, Beta, Theta and Gamma wave proportion of EEG in three groups of mice in different states **G**. Schematic Diagram of Mouse Cortical EEG Recording and optical fiber **\*\*\*p<0.001, **p<0.01, *p<0.05, PS photostimulation**

For Delta and Theta wave, the average proportion in the three groups during anesthesia had no significant difference (p>0.05, Fig 6 B and 6 E). As for Alpha wave, the average proportion of Alpha wave in vehicle group during maintenance phase was significantly higher than that in the recovery phase (P<0.05). And the average proportion of Alpha wave in PS (15mW, 10min) and PS (15mW, 15min) during maintenance phase was significantly higher than that in the recovery phase (P<0.05, P<0.05, Fig 7 C). For Beta wave, the vehicle group in the awake state was significantly lower than that in the anesthesia state (P<0.05, Figure 7 D). As for Gamma wave, the Gamma wave proportion in vehicle group was significantly higher in the awake stage than that in maintenance (P<0.05) and recovery (P<0.05). In the group with PS (15mW, 10min), the Gamma wave proportion was significantly higher in the recovery than that in maintenance (P<0.05). In the group with PS (15mW, 15min), the Gamma wave proportion was significantly lower in the maintenance than that in wakefulness (P<0.05) and recovery stage (P<0.01, Fig 7 F). These indicates EEG activity alters in different stages, especially Gamma wave.

### 3.7 Effect of ICV 5-HT 1A receptor antagonist on Induction time and Emergency time of sevoflurane anesthesia

pAAV-TPH2 PRO-ChETA-EYFP-WPRES-PAS was delivered into BLA 3 weeks before anesthesia, and a lateral ventricle cannula as well as an optical fiber were implanted. The mice were divided into four groups: vehicle + no PS, 1A(-) + no PS, vehicle + PS, 1A(-) + PS. The results showed the administration of 5-HT 1A receptor antagonist and photostimulation 5-HT neuron in BLA had no significant effect on the induction time of sevoflurane (P>0.05, Fig 8 B).

**Figure 8.**
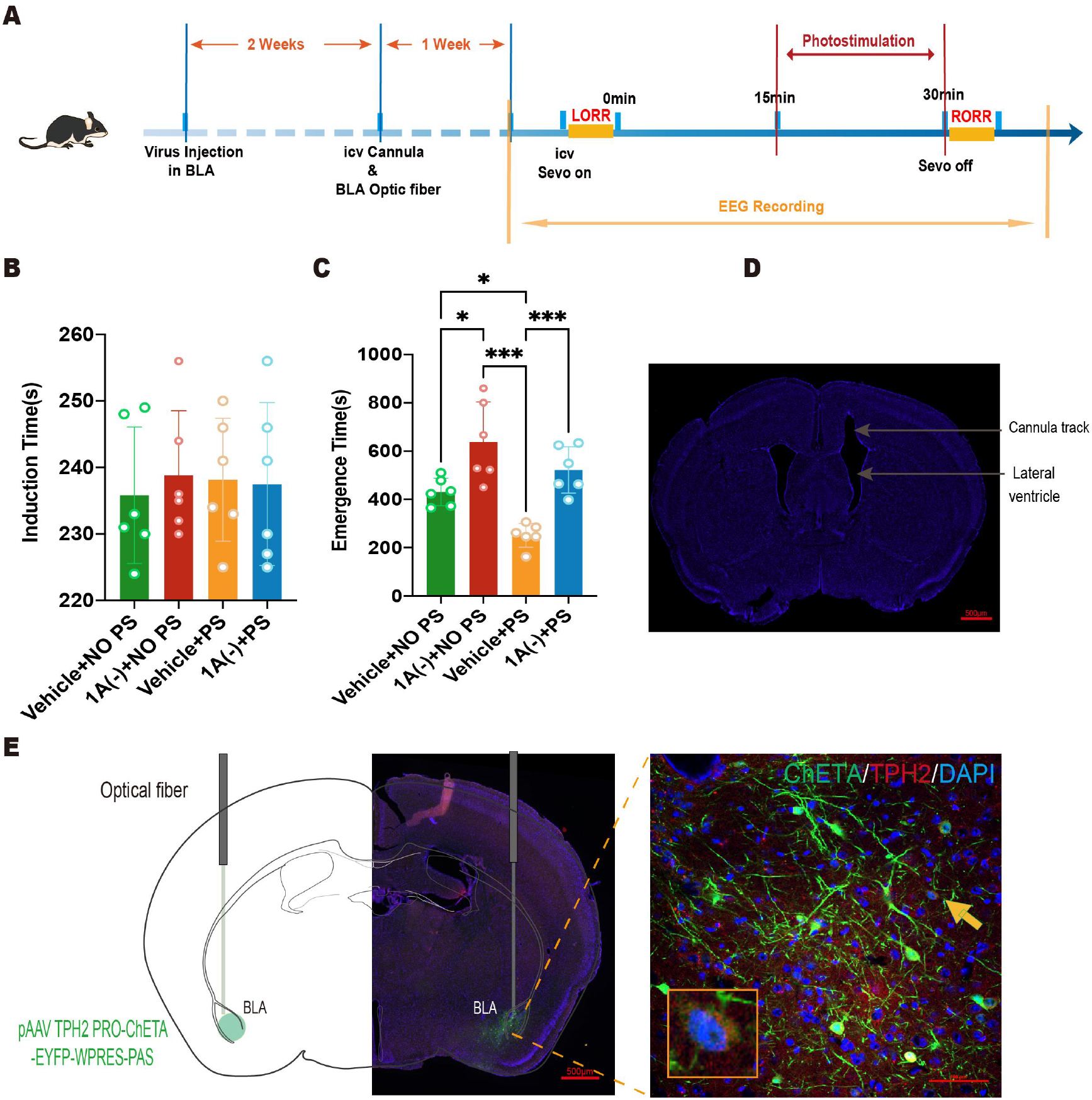
The suppressive effect of ICV 5-HT 1AR on Induce time and Emergency time. **A**. Experimental protocol for lateral ventricle administration and optogenetic stimulation of BLA 5-HT neurons. **B-C**. The effect of 5-HT 1AR and BLA 5-HT neuron photostimulation on Induce time and Emergency time of sevoflurane anesthesia **D**. A representative photomicrograph shows the locations of ICV cannula and optic fiber. **E**. A representative photomicrograph shows the locations of optical fiber and virus expression. 1A (-)= WAY 100635; **\*\*\*p<0.001, **p<0.01, *p<0.05, PS=photostimulation, ICV= intracerebroventricular**

As for the Emergency time, the group treated with 1A(-) + no PS was significantly longer than these treated with vehicle + no PS (P<0.05) and vehicle + PS (P<0.001), indicating 5-HT1AR was involved in mediating the recovery of sevoflurane anesthesia. And the group treated with 1A(-) + PS had significant longer Emergency time than the group received vehicle + PS, which suggested the suppressive effect of 5-HT 1AR could reverse the positive effect of activation 5-HT neurons in BLA.

### 3.8 The effect of ICV 5-HT 1A receptor antagonist on EEG activity

rAAV-TPH2-GCaMP6S-WPRE-hGH was delivered into BLA 3 weeks before anesthesia, an optical fiber, and an ICV cannula microinjecting 5-HT 1A(-) with a headstage for EEG was implanted 1 week before experiments. The mice were divided into four groups: vehicle + no PS, vehicle + PS, 1A(-) + no PS and 1A(-) + PS. The activity of cortical EEG in each group decreased significantly after LORR, and kept at a low level during the maintenance of anesthesia (Fig 9 A).

**Figure 9.**
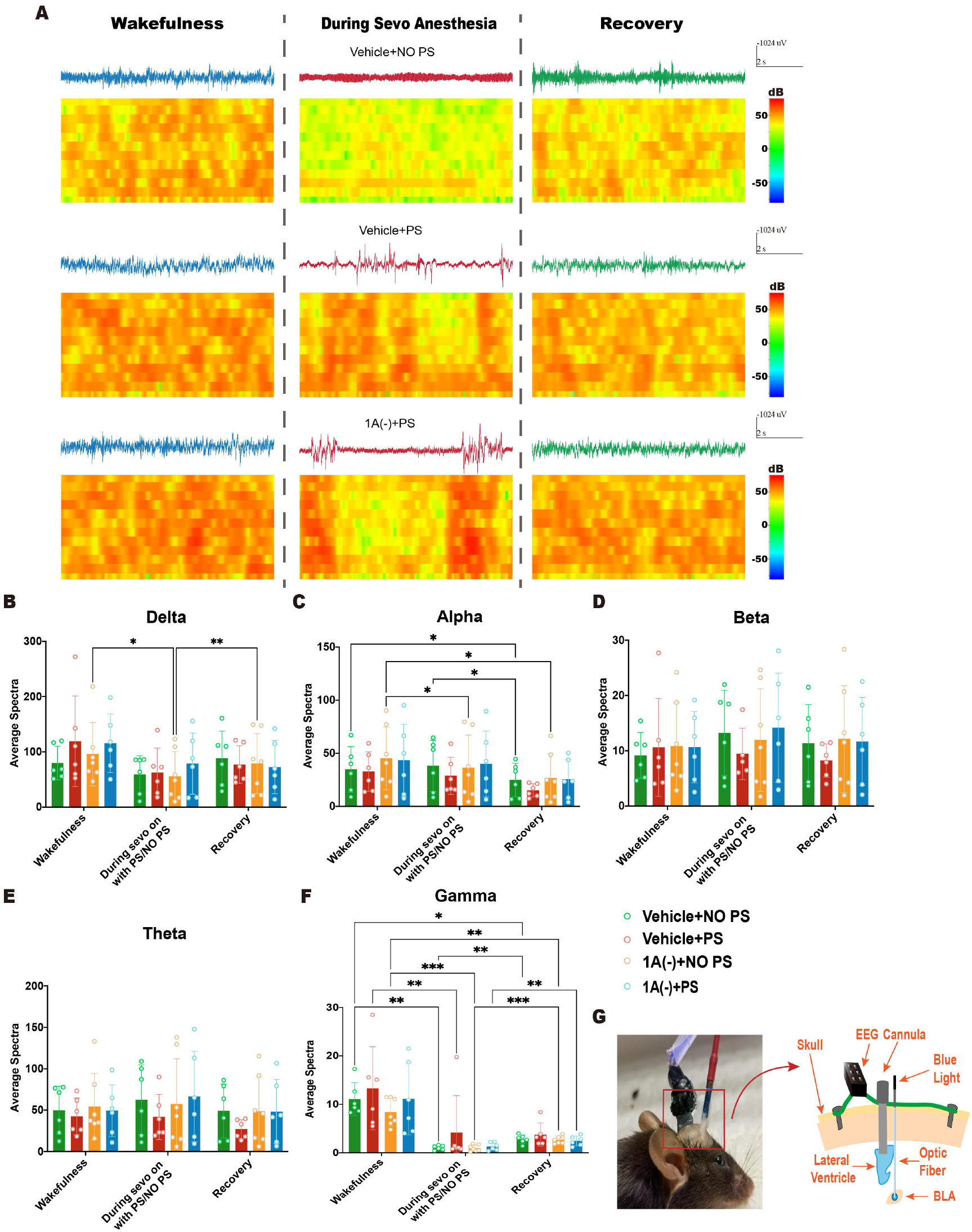
Effect of ICV 5-HT 1AR and photogenetic activation of 5-HT neurons in BLA on EEG activity. **A**. EEG and spectrum of mice in vehicle group + no PS, vehicle + PS, 1A(-) + no PS and 1A(-) + PS under awake, anesthetized and recovery states. **B-F**. Delta, Alpha, Beta, Theta and Gamma wave proportion of EEG in four groups of mice in different states **G**. Schematic Diagram of Mouse Cortical EEG Recording, ICV cannula and optical fiber. 1A (-)= WAY 100635; **\*\*\*p<0.001, **p<0.01, *p<0.05, PS=photostimulation, ICV= intracerebroventricular**

For Beta and Theta wave, the average proportion in the four groups during anesthesia had no significant difference (p>0.05, Fig 9 D and 9 E). As for Delta wave, the average proportion of Delta wave in 1A(-) + no PS group during maintenance phase was significantly lower than that in the awakening phase (P<0.05) and recovery phase (P<0.05, Figure 9 B), suggesting 5-HT 1A receptor may involve in mediating EEG activity during sevoflurane anesthesia. And the average proportion of Alpha wave in 1A(-) + no PS group during maintenance phase was significantly lower than that in the wakefulness phase (P<0.05, Fig9 C).

As for Gamma wave, the Gamma wave proportion in 1A(-) + no PS group was statistically lower in the anesthesia stage than that in awake (P<0.001) and recovery state (P<0.001) and the EEG activity did not return to the same level after RORR (P<0.001). In the group with 1A(-) + PS, the Gamma wave proportion was significantly higher in the recovery than that in maintenance (P<0.01, Fig 9 F). These indicates 5-HT 1A receptor in BLA may participate in mediating EEG activity during sevoflurane anesthesia, especially Gamma wave.

### 3.9 5-HT neuron projection from DR to BLA was established by the application of the nerve tracer CTB-555

To further retrograde the 5-HT neuron projection in BLA, we microinjected CTB-555 in BLA and then perfused after 1 week. Immunohistochemical results suggested that DR 5-HT neuron might project to BLA (FIG 10).

**Figure 10.**
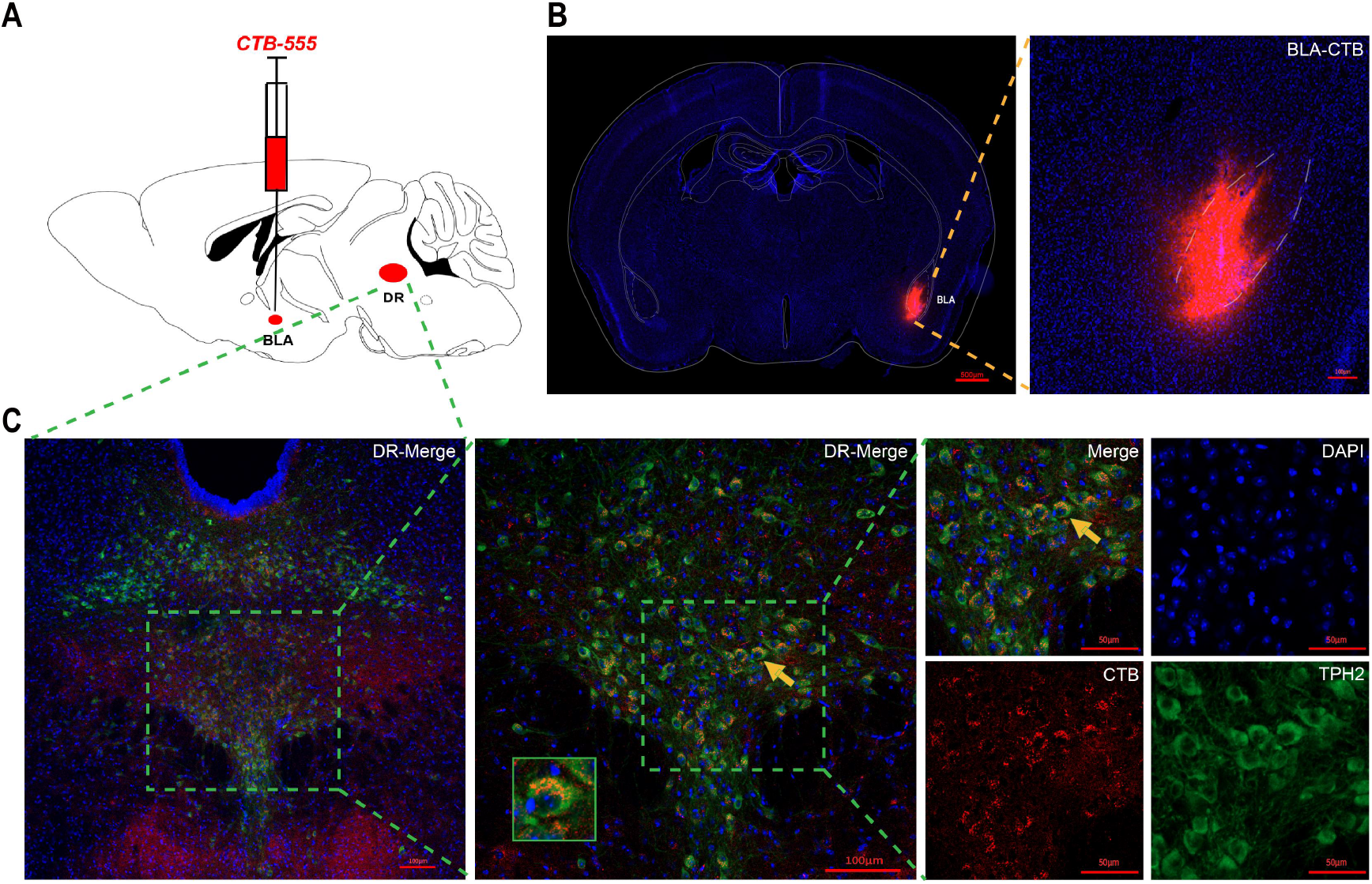
Neural projection from DR to BLA was established by microinjecting nerve tracer CTB-555 in BLA. **A-B**. Representative coronal brain slice, showing the location of CTB-555 injected in BLA. **C**. Projection from DR to BLA with co-expression of CTB-555, TPH2 and DAPI.

### 3.10 5-HT neural projection from DR to BLA involved in reducing the Emergency time of sevoflurane anesthesia

pAAV-TPH2 PRO-ChETA-EYFP-WPRES-PAS was delivered into DR 3 weeks before anesthesia, and optical fibers were implanted 1 week before experiments (Figure 11A). We selectively activated 5-HT neuron in DR by applying photostimulation.

**Figure 11.**
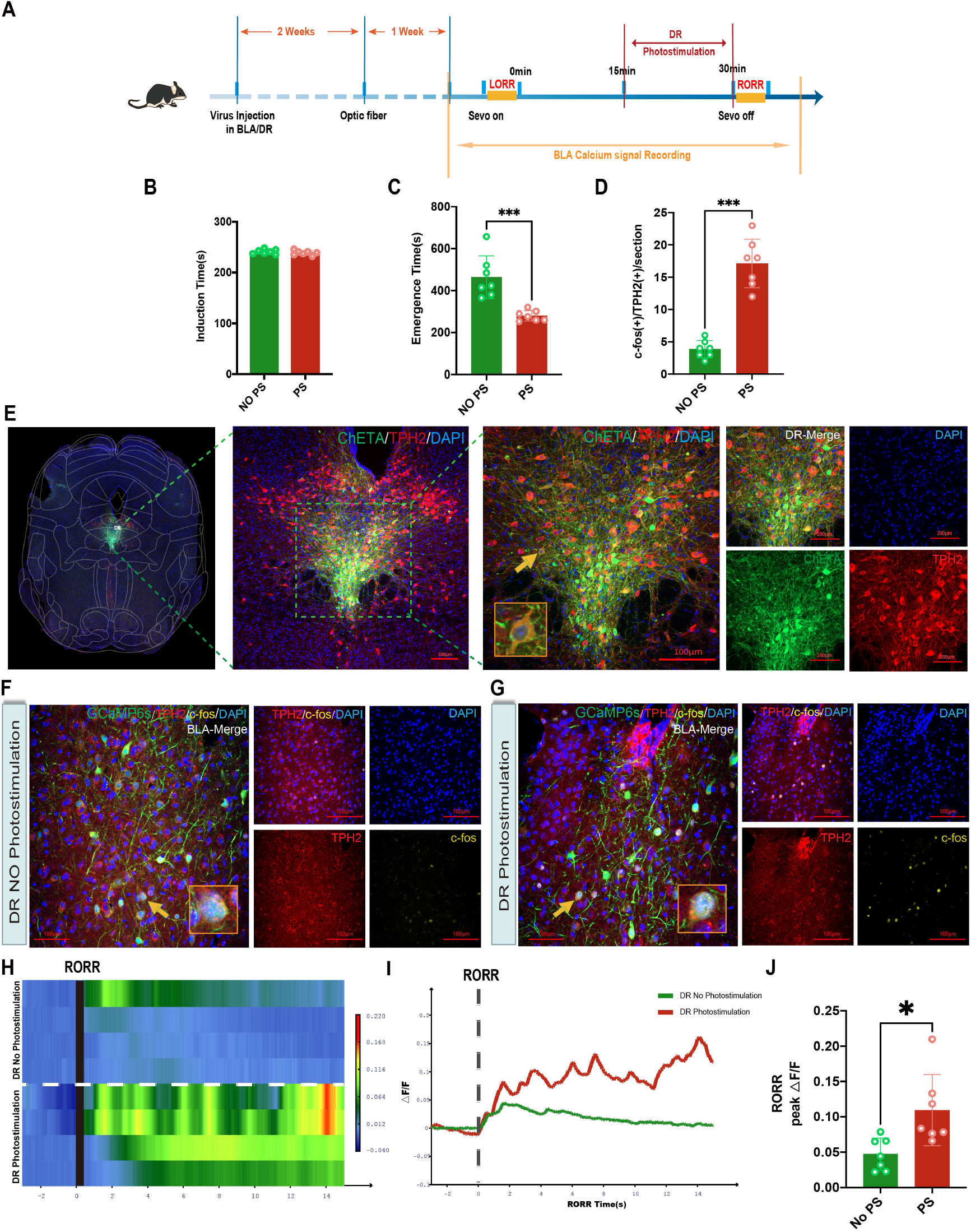
Photostimulation of DR 5-HT neuron influenced the expression of c-fos in BLA and calcium signaling in BLA 5-HT neuron during RORR. **A**. Experimental protocol for BLA 5-HT neuron photometric recordings and optogenetic stimulation of DR 5-HT neurons. **B**. The effect of activating DR 5-HT neuron on the Induce time, Emergency time during sevoflurane anesthesia and the expression of c-fos in BLA **C**. Shown are representative images of the tracks of optic fibers implanted into DR and staining for ChETA, TPH2 and DAPI in the DR. **D**. Representative images staining for c-fos, TPH2 and DAPI in DR **E**. The photometry recordings of the group with DR PS and without DR PS during the period of RORR **F-G**. There was significant difference in peak ΔF/F of BLA between DR with PS and DR with PS. **\*p<0.05, PS= photostimulation**

Induction time in control group without PS and group with PS showed no statistical difference. Compared with the control group without PS, there was significant difference in the Emergency time of mice in experimental group with PS (P<0.001). Quantification of c-fos(+)/TPH2(+) cells in BLA with PS in DR were significantly more than these cells in BLA without PS in DR (p<0.05, Figure 11B).

Calcium signaling within neurons of bilateral BLA was recorded by photometry in mice infected with GCaMP6s in the group treated with or without photostimulation in DR during maintenance stage (Figure 11 E). We observed obvious difference in peak ΔF/F in BLA between the control group and photostimulation group (Fig 10F). Compared with the vehicle group, peak ΔF/F of RORR was significantly increased in the photostimulation group (p<0.05, Fig 11 G). These data indicate that 5-HT neural projection from DR to BLA involved in reducing the Emergency time of sevoflurane anesthesia.

## Discussion

The researches on the specific mechanism of arousal from general anesthesia, and the mechanism of reversible loss of consciousness during general anesthesia is the focus of the field of anesthesia. In this study, we used optogenetic techniques and intraperitoneal injection and intracerebroventricular injection to reveal the role of BLA in emergence from general anesthesia. And the effects of anesthesia and excited the serotonin neurons in BLA on the electroencephalogram (EEG) of the mice were observed by cortical EEG recording. The present results demonstrated that the stimulation of BLA could reduce the emergency time from general anesthesia, which might through 5-HT neurons, particularly in the DR.

General anesthetics have been used for more than 170 years, which can cause loss of consciousness and analgesia, and ensure the success of surgery. However, the mechanism of general anesthesia is still unclear. The journal Science listed “How general anesthetics work” as one of 125 frontier questions in science. Using D1R-Cre mice, the study found that dopamine D1R-positive neurons showed decreased activity during sevoflurane anesthesia induction and increased activity during emergence by in vivo fiber calcium signals and EEG-EMG recordings. Blue light specifically activates D1R positive neurons in the nucleus accumbens of mice under continuous stable anesthesia, which can rapidly induce cerebral cortex activation and behavioral awakening^27^.

It suggests that the mechanism of general anesthetics may intersect with the sleep-wake regulatory neural circuit. The neural networks, involved in the mechanism of general anesthetic agents and sleep-wake, are comprised of the basal forebrain, the preoptic area of the hypothalamus which contains the ventrolateral preoptic (VLPO) nucleus, the posterolateral region of the hypothalamus, the brainstem, and the thalamus^28^. The lateral hypothalamic area (LHA) contains wake-promoting orexinergic (hypocretin) neurons, which is essential for stabilizing the sleep-wake state, activing during wakeful and REM sleep, and capable of sending projections throughout the brain, including to the VLPO, to maintain wakeful state ^29, 30, 31^. The prefrontal cortex (PFC) receives input fibers from the dorsomedial thalamus and has extensive connections with subcortical wakeing-related nuclei, including the serotonergic dorsal raphe nucleus (DR), the noradrenergic locus ceruleus (LC), the cholinergic basal forebrain, and the dopaminergic ventral dorsal tegmental area (VTA). Studies have shown that the inactivation of the PFC can delay the emergence from general anesthesia in rats, and the PFC also plays an important role in awakening. However, it should be noted that the role of the posterior cortex in awakening has also been confirmed in many other experiments^32^.

BLA is associated with many behaviors, such as fear, aggression, learning, and memory ^33^. From the amygdala (via the central nucleus) there is a massive projection to the brainstem regions involved in the arousal and REM waves and ponto-geniculo-occipital (PGO) waves ^34, 35^. The BLA receives cortical and subcortical inputs and projects to brainstem, hypothalamus, forebrain, and cortical regions associated with hormonal, autonomic, and behavioral aspects of stress responses and is also involved in the sleep-wake cycle ^36, 37^. There are still few studies on the role of BLA in sleep-wake and general anesthesia mechanism, so we chose BLA as the target to explore the mechanism of emergence from general anesthesia.

5-HT neurons are widely distributed and projected in the central nervous system, and their physiological functions are complex due to their numerous receptor subtypes. In the study of sleep-wake regulation drugs, it is found that most drugs act through different subtypes of 5-HT receptor. Studies have shown that 5-HT in the hippocampus has a sleep-promoting effect by changing the content of 5-HT in the thalamic reticular nucleus and activating or inhibiting 5-HT1A receptors to produce sleep-promoting or wake-promoting effects. Microinjection of 5-HT1A receptor agonist into the DRN stimulated the autoreceptor of 5-HT1A, and increased REMS were found in cats and rats ^38^. Injection of p-MMPI, a 5-HT1A receptor antagonist, into the DRN resulted in a reduction in REMS and spontaneous firing of DRN neurons. The 5-HT1B receptor in the medial preoptic nucleus also has a role in arousal. Studies have found that injection of 5-HT3 receptor agonist in DRN can reduce REMS and SWS and promote arousal^39^. However, injection of 5-HT6 receptor antagonist had no significant effect on REMS. After treatment with 5-HT7 receptor antagonist or knockdown of 5-HT7 gene, REM sleep time was reduced in rats, which was later found to be indirectly mediated by 5-HT1A receptor. Our previous study showed that activation of 5-HT 1A receptor by lateral ventricle and DR intranuclear injection can significantly shorten the emergence time of sevoflurane anesthesia. Earlier studies have suggested that 5-HT is a ‘sleep’ neurotransmitter as the lesion of the raphe nuclei leads to insomnia in cats^40^. The previous studies have shown that the extracellular level of 5-HT in BLA is lower during sleep than in waking which might be mediated by the serotonergic projection of the DRN to the BLA ^41, 42^. Therefore, we investigated the relationship between BLA and 5-HT in DR during emergence from general anesthesia, and whether 5-HT 1A receptors play a major role in reducing the emergence time.

In addition to the role of 5-HT system in BLA regulation of emergence time of general anesthesia, norepinephrine (NE), synthesized by the neurons of brainstem locus ceruleus (LC) nuclei, may also be involved during the arousal in BLA. NE-containing LC neurons are interconnected with forebrain structures, including the amygdala, and has been implicated in the regulation of the sleep-wake cycle^43, 44, 45^. It has been shown that extracellular levels of NE in the LC and amygdala gradually decrease from wakefulness to slow wave sleep to REM sleep ^46^. Therefore, the BLA may receive information from different neurotransmitters and brain regions, except 5-HT in DR, for integration at the same time. Further studies on the connections and projections between the BLA and other brain regions during the emergency of general anesthesia will be conducted in the future.

## Figure Legends

**Table 1.**
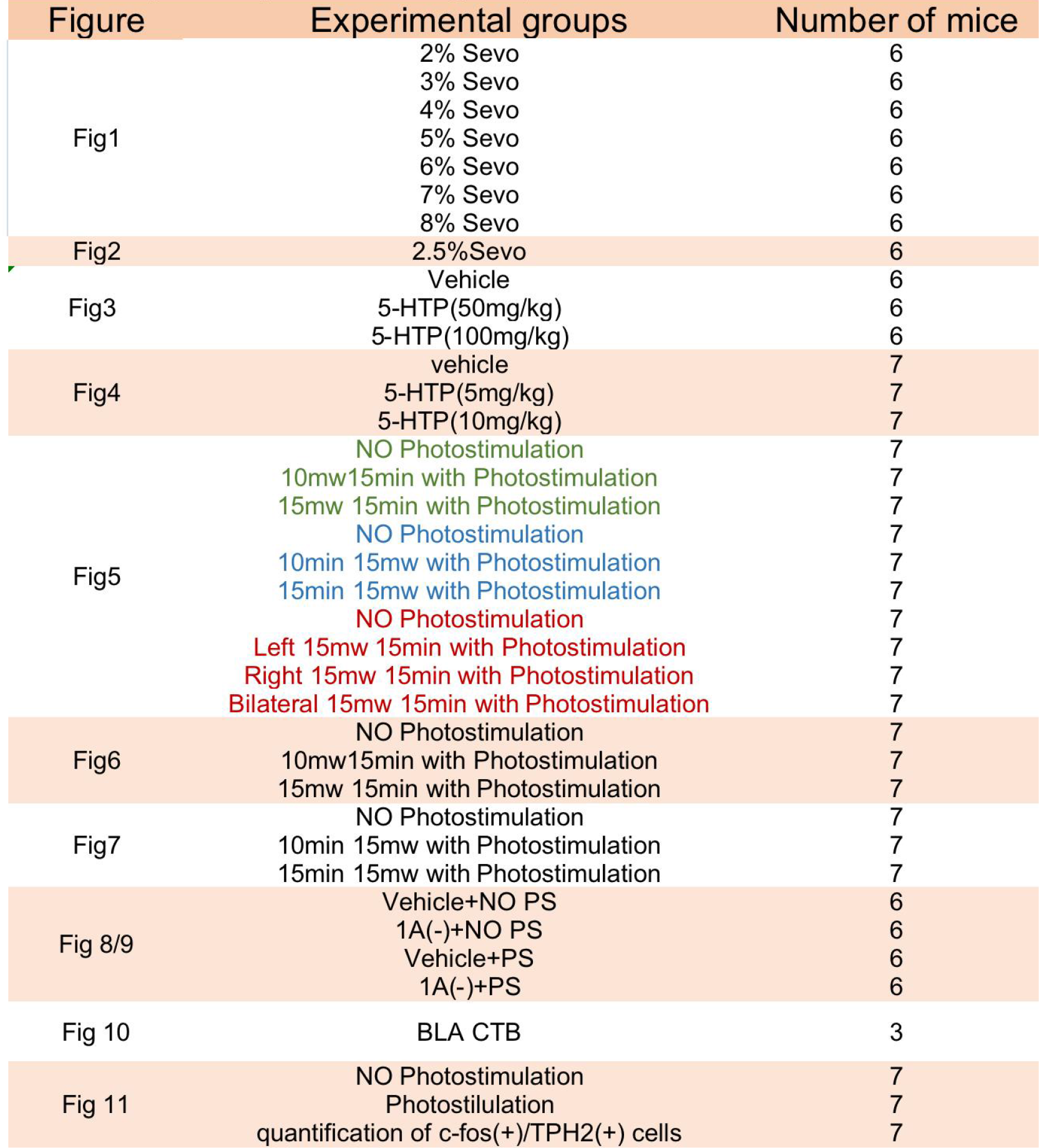
summary of experimental groups of C57BL/6J mice

**Table 2.**
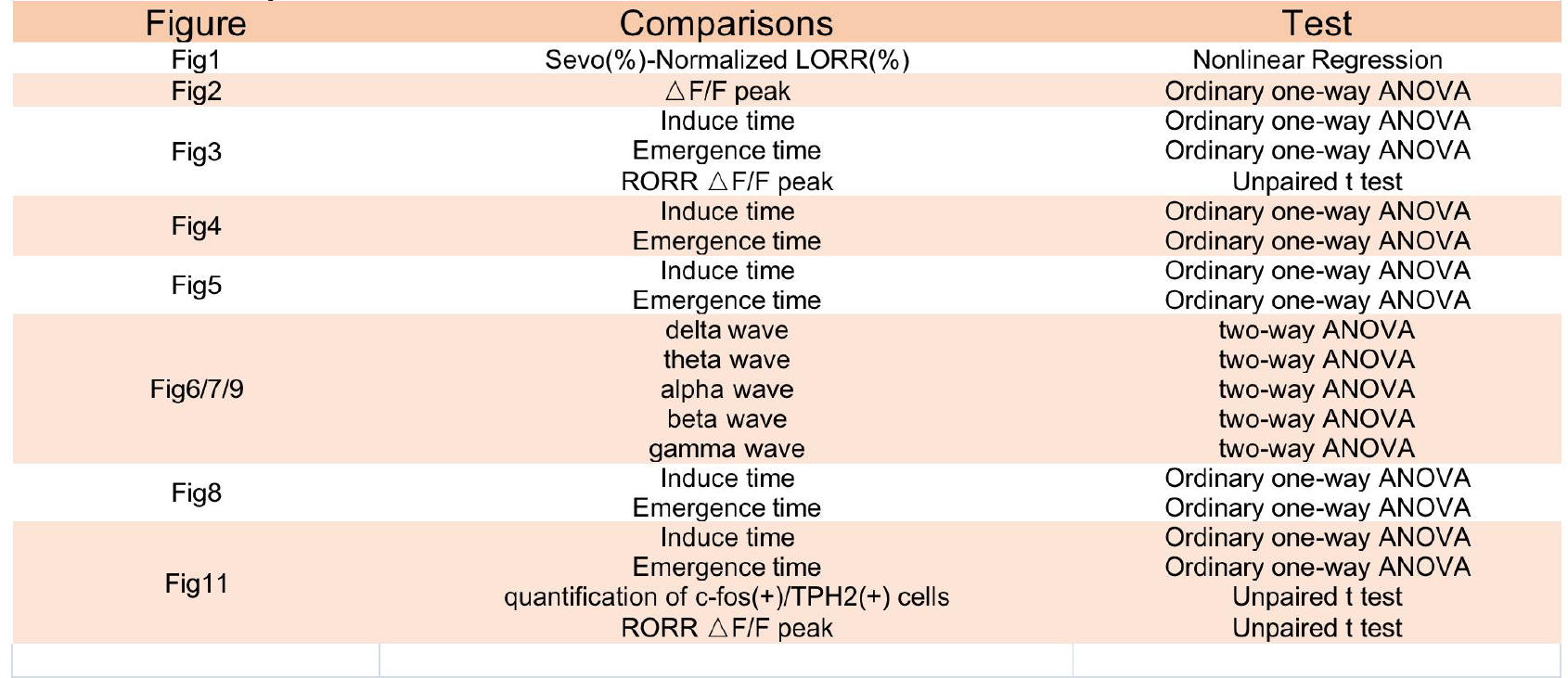
statistical analysis

## Acknowledgments

The work was supported by the National Natural Science Foundation of China (Grant.NO: 81771403, 81974205); by the Natural Science Foundation of Zhejiang Province (LZ20H090001); by the Program of New Century 131 outstanding young talent plan top-level of Hang Zhou to HHZ

**All authors declare no competing interests**.

